# Uncovering indirect interactions in bipartite ecological networks

**DOI:** 10.1101/315010

**Authors:** Benno I. Simmons, Alyssa R. Cirtwill, Nick J. Baker, Lynn V. Dicks, Daniel B. Stouffer, William J. Sutherland

## Abstract

Indirect interactions play an essential role in governing population, community and coevolutionary dynamics across a diverse range of ecological communities. Such communities are widely represented as bipartite networks: graphs depicting interactions between two groups of species, such as plants and pollinators or hosts and parasites. For over thirty years, studies have used indices, such as connectance and species degree, to characterise the structure of these networks and the roles of their constituent species. However, compressing a complex network into a single metric necessarily discards large amounts of information about indirect interactions. Given the large literature demonstrating the importance and ubiquity of indirect effects, many studies of network structure are likely missing a substantial piece of the ecological puzzle. Here we use the emerging concept of bipartite motifs to outline a new framework for bipartite networks that incorporates indirect interactions. While this framework is a significant departure from the current way of thinking about networks, we show that this shift is supported by quantitative analyses of simulated and empirical data. We use simulations to show how consideration of indirect interactions can highlight ecologically important differences missed by the current index paradigm. We extend this finding to empirical plant-pollinator communities, showing how two bee species, with similar direct interactions, differ in how specialised their competitors are. These examples underscore the need for a new paradigm for bipartite ecological networks: one incorporating indirect interactions.

## Introduction

Ecological communities are widely represented as bipartite networks that depict interactions between two groups of species, such as plants and pollinators. These networks are used to answer a diverse range of questions about community structure, such as whether antagonistic and mutualistic communities have different architectures (Fontaine et al. 2011, Morris et al. 2014); how plant-frugivore communities at forest edges differ from those in forest interiors (Menke et al. 2012); whether fluctuations in species and interactions over time alter network structure (Petanidou et al. 2008); and whether individual pollinators vary in their use of floral patches (Dupont et al. 2014).

For over thirty years, the framework for characterising the structure of bipartite networks has remained unchanged: indices, such as nestedness and species degree, are used to describe either whole-network topology or the roles of individual species with a single summary statistic. However, while these indices have greatly improved our understanding of community structure, they also suffer from a substantial, but largely ignored, ecological limitation: reducing a complex network to a handful of one-dimensional metrics necessarily involves a loss of information. This is because indices are insensitive to changes in pairwise species interactions: different network configurations can have identical index values (Olito and Fox 2015). Often this means discarding important detail about indirect interactions. For example, let there be two communities: in the first community, plant *i* is pollinated by one species, *j*; in the second community, *i* is still only pollinated by *j*, but *j* also pollinates plants *k*, *l* and *m*. We cannot distinguish the two situations by examining, for example, the degree of *i* because degree discards all information on indirect interactions: we know that *i* has a direct interaction with *j*, but we do not know whether *j* is an obligate specialist on *i* or a generalist visiting several other plants.

This loss of ecological detail resulting from the use of network indices is concerning as it puts many studies describing network structure directly at odds with a large literature that has repeatedly documented important and widespread indirect effects in nature (Wootton 2002). For example, in mutualistic networks, dynamical models (which use the whole network as the skeleton of dynamics and therefore incorporate indirect interactions) have shown that indirect effects are a major process governing coevolution (Guimarães et al. 2017), while in host-parasitoid communities, apparent competition and even apparent mutualism can occur when herbivorous insects influence each other through shared natural enemies (Morris et al. 2004, Frank van Veen et al. 2006, Tack et al. 2011). Similarly, indirect effects between co-flowering plant species in pollinator communities can range from facilitation, where the presence of one plant increases the frequency of pollinator visits to another, to competition, where one plant attracts pollinators away from another (Mitchell et al. 2009, Morales and Traveset 2009, Carvalheiro et al. 2014). Indirect interactions are therefore a fundamental component of ecosystems, driving ecological and evolutionary processes to an equal, or greater, extent than direct interactions (Vandermeer et al. 1985, Strauss 1991, Bailey and Whitham 2007, Martínez et al. 2014, Guimarães et al. 2017). Widespread, uncritical use of network indices risks missing all or part of this component.

Here we advocate a new way of thinking about bipartite networks that moves beyond network- and species-level structural indices to also incorporate indirect interactions. We argue for conceptualising networks as a collection of constituent parts or ‘building blocks’ using the emerging concept of bipartite motifs (subgraphs representing patterns of interactions between a small number of species). We outline the theory, applications and future directions of this framework. Although what we propose is a profound shift in the way we conceptualise bipartite networks, we show that it is well supported by simulated and empirical data, using three analyses to demonstrate the importance of indirect interactions and how they are captured by the proposed framework. First, we use three six-species networks to show that indirect interactions are necessary to accurately describe a species’ role in even a small community. Through simulation, we then generalise this finding to a large ensemble of networks with diverse sizes and structures to establish and quantify how communities with similar overall properties can exhibit remarkable dissimilarity in their indirect interaction structure. Finally, we demonstrate these results in an empirical context, highlighting how indirect interactions can result in ecologically important differences between two pollinator species with similar direct interactions. We also assess the robustness of the framework to sampling effort and propose several hypotheses about how our understanding of ecological communities might change if indirect interactions were incorporated. We hope to motivate uptake of a new research agenda incorporating indirect effects in both empirical and theoretical studies of ecological networks.

## Indirect interactions and the index paradigm

Consider the role of species *B* in the three communities shown in Fig. 1a. Based only on its direct interactions, the role of *B* is identical in all three communities: *B* interacts with *E*. However, by considering the interactions of *B*’s partner *E*, *B*’s role in networks I and III can be distinguished from its role in network II: in networks I and III, *B* competes with *A* and *C* for the shared resource *E*, while in network II, *B* competes with only *A*. Furthermore, by considering the interactions of *A* and *C* (*B*’s partners’ partners), the roles of *B* in networks I and III can also be distinguished: in network I, *B*’s competitor *C* is a specialist on resource *E*, while in network III, *C* also visits *F*. Similarly, while *A* is a super-generalist in network I, visiting every resource in the community, in network III it has a narrower diet breadth, visiting only *D* and *E*.

**Figure 1:**
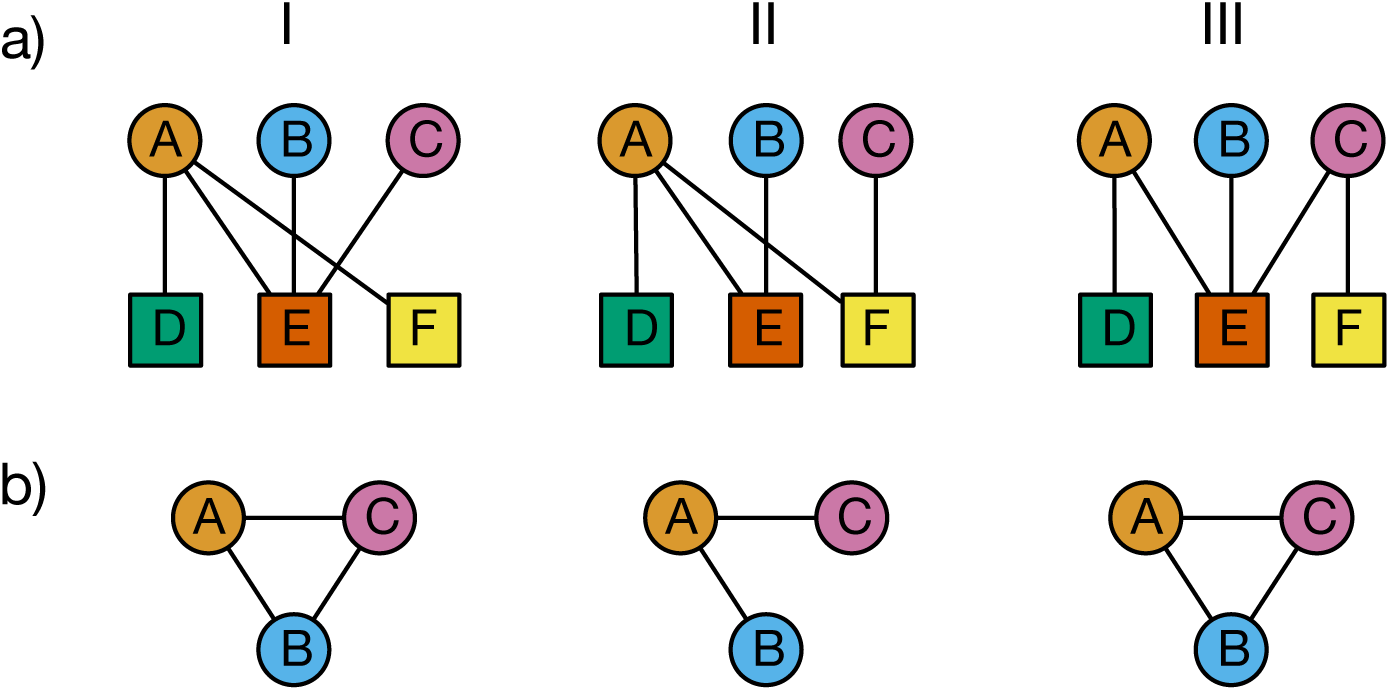
Three example networks (a) and the corresponding one-mode projections for species in the upper level (b).

This simple example shows how indirect interactions are necessary to give a complete picture of a species’ role, even in a small community: between the three networks, *B* differs both in the number of competitors it has and in how specialised these competitors are on the shared resource *E*. Such differences are likely to have important ecological consequences. To capture this detail, it was not sufficient to consider only *B*’s direct interactions, or even the interactions of *B*’s partner; rather we had to go ‘deeper’ and consider the interactions of *B*’s partners’ partners to differentiate its role in all three networks.

Many indices that capture interaction patterns at the level of individual species – such as degree, dependence (the strength of an interaction between species *i* and *j* as a proportion of *i*’s total interaction strength) or species strength (sum of dependencies on a species) – are largely based on direct interactions and so do not give a complete picture of *B*’s role in the three communities. Other species-level indices, such as *z*- and *c*- scores, do consider indirect interactions, but only with respect to modules (groups of species connected more to each other than to other species in the network). *z*-scores quantify a species’ connectivity within modules, while *c*-scores (also known as the participation coefficient) quantify a species connectivity among modules. In all three networks, *B* has identical *c*-scores, while *z*- scores for *B* are identical in networks I and III. Various centrality indices also incorporate indirect interactions, but rely on the one-mode projection of the bipartite network, where species in one set are linked when they share one or more partners in the other set (Jordán et al. 2007). As can be seen in Fig. 1b, this compression necessarily leads to a substantial loss of information (Zhou et al. 2007, Saracco et al. 2017). For example, interactions with specialist species such as *D* are not considered as, by definition, specialists only interact with one species. Consequently, betweenness centrality (the number of shortest paths between two species passing through a focal species) and closeness centrality (the mean shortest path between the focal species and all other species) values for *B* are identical in all three networks. Finally, even multivariate combinations of common indices describing whole-network structure cannot distinguish between these three situations because all three communities have identical connectance, nestedness and modularity.

This is an example of the Goldilocks principle: by accounting for all interactions simultaneously, indices characterising whole-network patterns can be too coarse to detect fine differences. Conversely, by considering too little of the indirect interaction structure, indices describing individual species roles can miss differences beyond their local scope. In both cases, indirect interactions occurring at a level between these whole-network- and species-scales – that is, at the meso-scale – are missed. This problem arises from the limited amount of information a network index can provide: compressing a complex network into a single real number means that indices are only able to capture specific aspects of network structure or a species’ role rather than a complete picture of how each species is embedded in the community. Given the importance and ubiquity of indirect interactions there is therefore a need to move ‘beyond indices’ and adopt a new framework for describing network structure that uncovers the indirect interactions present in the meso-scale topology of networks.

## A framework for indirect interactions

We start by recognising the fact that any given network made up of *S* species can be broken down into a series of smaller subnetworks containing *n* species (where *n* < *S* and all species have at least one interaction). For example, network I in Fig. 1a includes five subnetworks containing two species (*A*–*D*, *A*–*E*, *A*–*F*, *B*–*E*, *C*–*E*) and six subnetworks containing three species (*D*–*A*–*E, D*–*A*–*F, E*–*A*–*F, A*–*E*–*B, A*–*E*–*C, B*–*E*–*C*). As there are a finite number of ways to arrange interactions between *n* species, there are also a finite number of possible subnetworks of size *n* that a network can contain. In other words, all bipartite networks, regardless of their complexity, are assembled from a limited number of parts or building blocks known as ‘bipartite motifs’ (Baker et al. 2015). For example, Fig. 2 shows all 44 possible motifs containing between two and six species. We argue that an understanding of these basic structural elements is required to capture the details of indirect interactions beyond the global and local features captured by network- and species-level indices (Milo et al. 2002).

**Figure 2:**
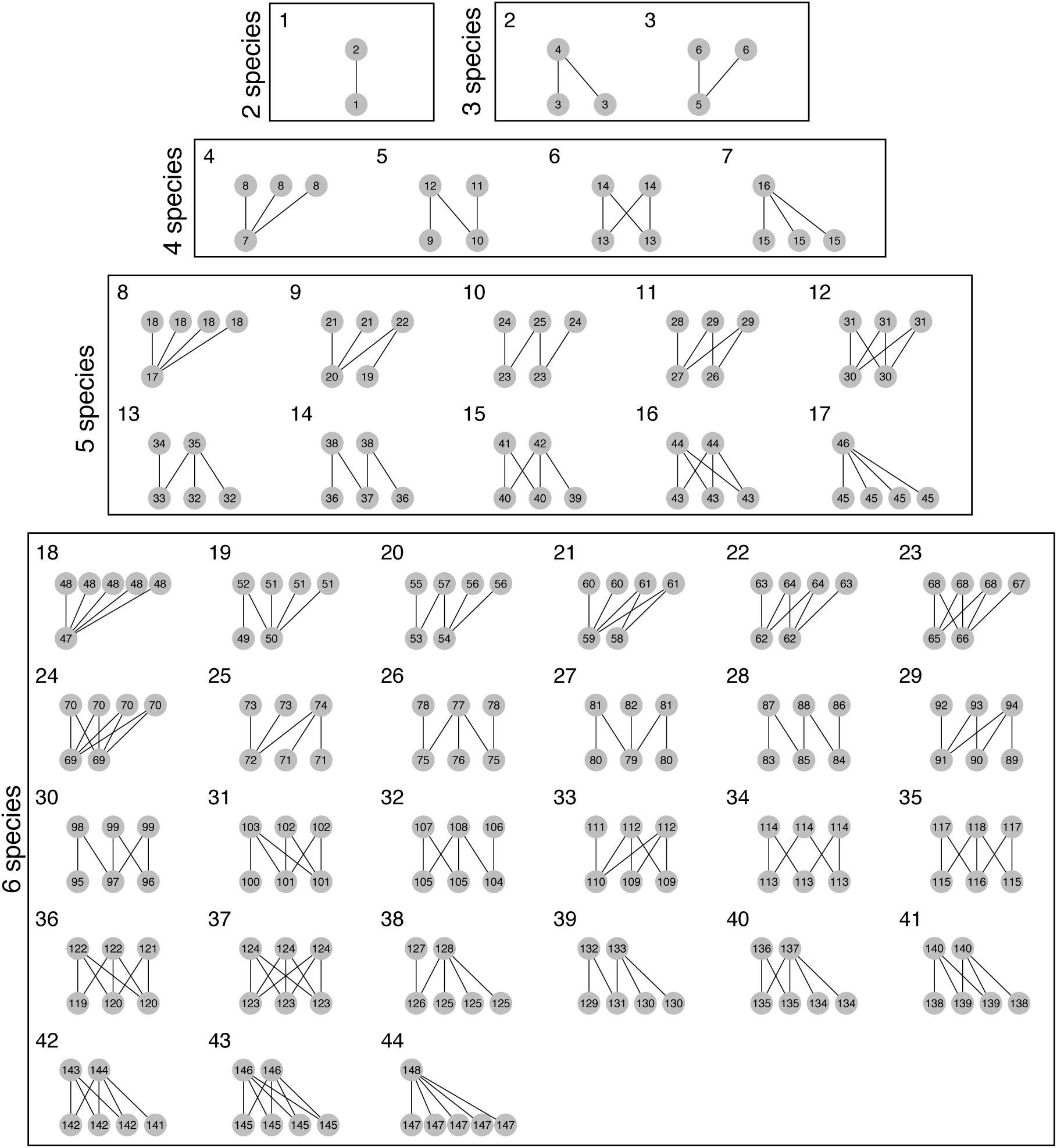
All possible two- to six-species bipartite motifs. Large numbers represent individual motifs. Small numbers within nodes represent unique position within motifs. In total there are 148 positions across 44 motifs.

To characterise network structure under this framework, networks are first decomposed into their constituent motifs, giving an inventory of the parts which make up the network. These simple lists show the frequency *c_i_* with which each motif *i* occurs in a network. This provides an *m*-dimensional ‘signature’ of a network’s structure, given by the vector 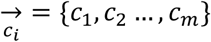, where *m* is the number of motifs counted. For example, Fig. 3 shows the constituent motifs of each example network from Fig. 1 and Fig. 4a shows the structural signature of each network. When viewed in this way, it becomes clear that each of these communities is made up of different parts, despite having similar or identical values of several common network indices.

**Figure 3:**
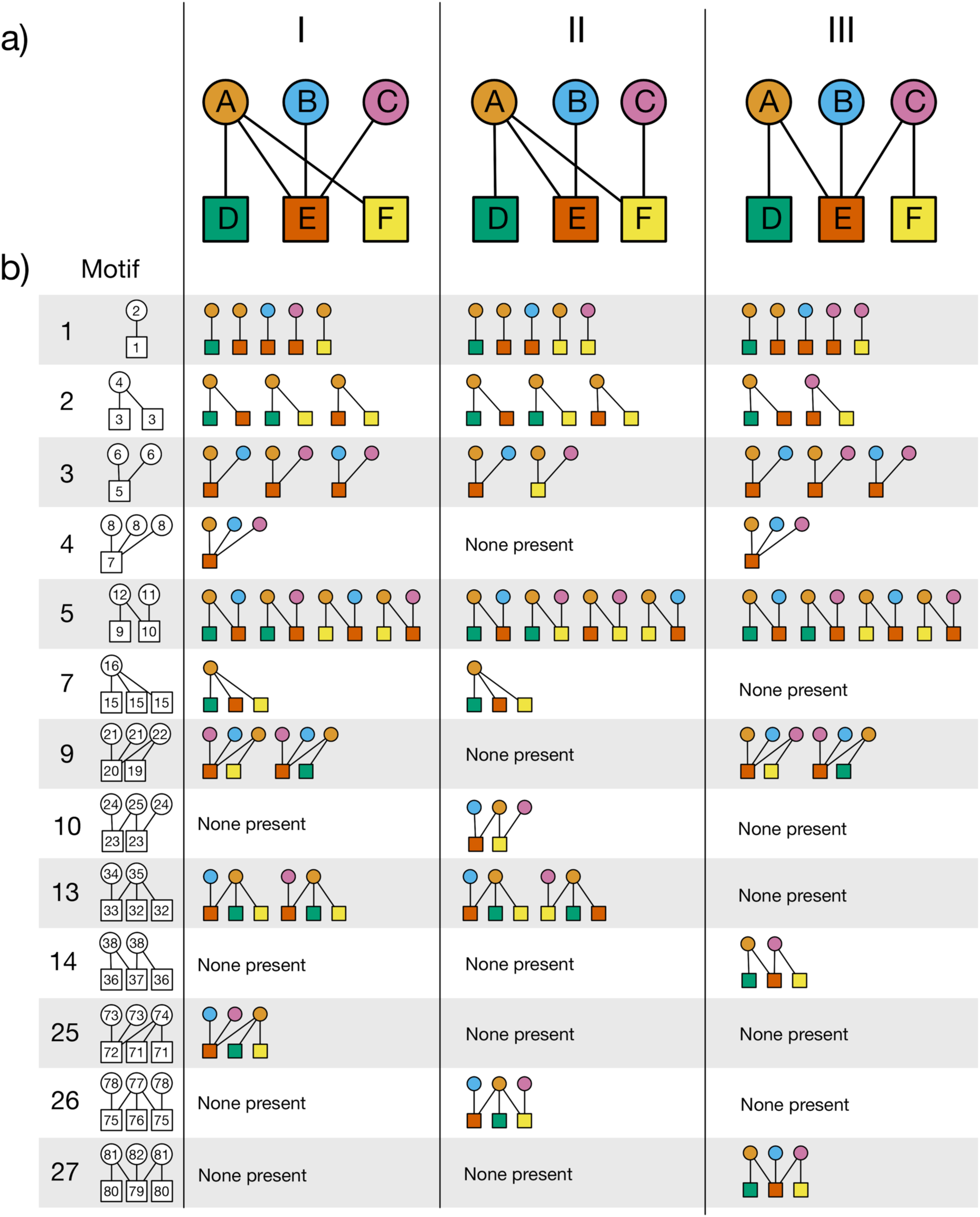
Decomposing three example networks into their constituent two-to six-node motifs. (a) Three example networks also used in Fig. 1a. (b) Table showing each network’s constituent motifs. The first column shows the motif being counted: the large number refers to the ID of the motif as given in Fig. 2; the small number within each node refers to the unique positions species can occupy within each motif as given in Fig. 2. The second, third and fourth columns show the occurrences of each motif in networks I, II and III respectively. Node colours refer to the species involved in each motif. For visualisation purposes, we exclude motifs which do not occur in any network, such as motif 8.

Within motifs, species can occupy different positions (Kashtan et al. 2004). For example, in motif five there are four unique positions, as each species interacts with a unique set of partners (Fig. 2). Considering all bipartite motifs up to six species, there are 148 unique positions (Fig. 2). Note that, due to symmetry, there may be fewer than *n* unique positions in a motif with *n* species. For example, in motif six there are only two unique positions, as both species in the top level interact with both species in the bottom level (Fig. 2). Therefore, a bipartite motif with *n* species can include between 2 and *n* unique positions. As these positions have distinct ecological meanings, a species’ role in a network can be defined by the frequency with which it occurs in each position (Stouffer et al. 2012, Baker et al. 2015, Cirtwill and Stouffer 2015). For example, Fig. 3b shows how, in network I, species *B* occurs once in position two, twice in position six, once in position eight, and so on. Generally, therefore, species roles are described by a vector 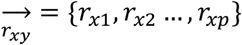, where *r_xy_* is the frequency with which species *x* occurs in position *y* and *p* is the number of positions counted. This vector can be thought of as a *p*-dimensional signature of a species’ role, or its multidimensional ‘interaction niche’. Fig. 4b shows the role signature of species *B* in the three networks from Fig. 1a. The roles are different in each network, demonstrating how this framework, by capturing indirect interactions at the meso-scale, distinguishes species roles that many indices cannot. Additionally, this did not require an *a priori* selection of the particular aspect of network structure to examine for differences; rather the approach is general, simply providing a detailed view of how *B* is embedded in the community.

**Figure 4:**
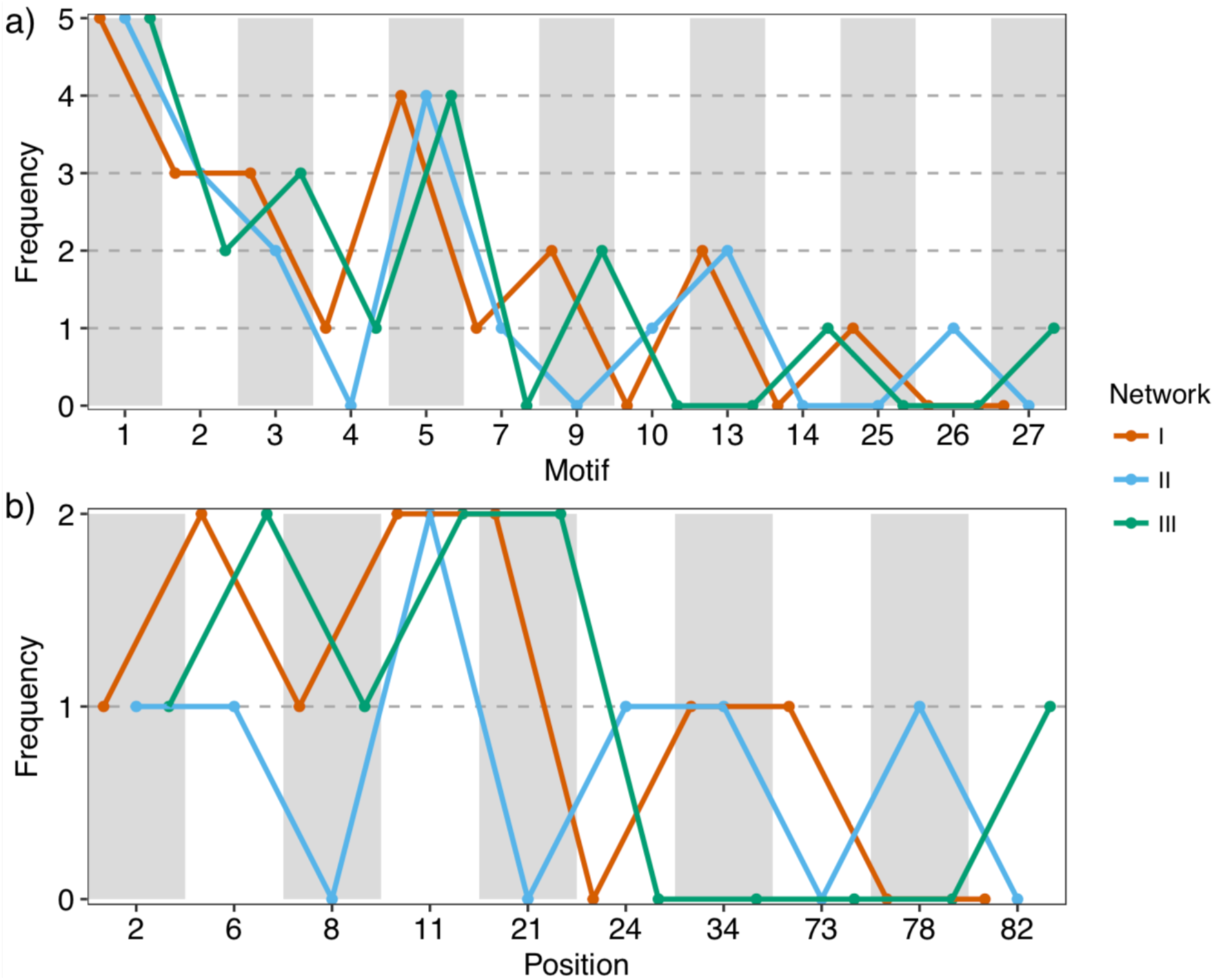
Motif and position frequencies of networks I, II and III from Fig. 1a and Fig. 3a. (a) The frequency with which each motif occurs in each network. For visualisation purposes, we exclude motifs which do not occur in any network. (b) The frequency with which species B occurs in each unique position within motifs in each network. For visualisation purposes, we exclude positions within which species B does not occur in any network. Motif and positions IDs correspond to those given in Fig. 2 and Fig. 3.

## Existing ecological research using bipartite motifs

To our knowledge, bipartite motifs have been used in only three published ecological studies. In one example, the motif role signatures of hosts and parasitoids were analysed in networks collected from forest patches in southern Finland over two years (Baker et al. 2015). The authors found that roles were significantly more similar within species than between species, suggesting that roles may be an intrinsic species property. Remarkably, species also exhibited this role fidelity between years, despite substantial species and interaction turnover. These findings may substantially improve our ability to predict changes in communities subject to environmental change.

In another example, motifs were used to disentangle the functional consequences of mutualism and antagonism in plant-animal interactions (Rodríguez-Rodríguez et al. 2017). The authors created an interaction typology of three-species motifs based on the type (mutualistic or antagonistic) and strength (weak or strong) of species interactions with *Isoplexis canariensis* in North-West Tenerife. The motifs were arranged along a mutualism-antagonism gradient, with plants interacting only with mutualists at one end, and antagonists at the other. Motifs in the middle of the gradient represented plants interacting with both mutualists and antagonists. Results showed that plant female reproductive success declined with increasing antagonism along the motif gradient, but that plants interacting with both mutualists and antagonists had greater importance in pollination networks, receiving and transferring more pollen. This study illustrates how plant reproductive success can be influenced by meso-scale variation captured by motifs.

Finally, community and interaction turnover was examined in a high-Arctic plant-pollinator network over fifteen years (Cirtwill et al. 2018). By calculating the roles of species using motifs, it was found that both plants and pollinator roles have changed significantly over time, potentially disrupting ecological functions. Given substantial climate change pressures in the Arctic, these findings suggest that the functioning of this network may be even more affected in the future. These three studies highlight the power of the motif framework for answering important ecological questions. However, the lack of studies using bipartite motifs suggests that many ecologists may be unaware of the approach or its benefits. In the examples that follow, for the first time, we explicitly quantify these benefits in simulated and empirical data, establishing that networks with similar overall properties can exhibit remarkable dissimilarity in their indirect interaction structure.

## Quantifying the loss of indirect interactions

In Figs. 1, 3 and 4, we used simple six-species networks to demonstrate how indices can mask potentially important meso-scale variation in indirect interactions. Here we generalise this effect to a large ensemble omf networks of varying sizes and structures using quantitative simulations. We first generated 20,000 bipartite networks containing 5 to 50 species in each set (giving 10 to 100 species) and with connectances ranging between the minimum required for each species to interact with at least one partner and 0.5. Networks were generated using the bipartite cooperation model (Saavedra et al. 2009). For each network, we characterised its structure at the macro-scale, using three whole-network indices (connectance, nestedness, modularity), and at the meso-scale, using the structural signature containing the frequencies of motifs up to five species. Nestedness was measured as NODF (Almeida-Neto et al. 2008) and modularity was calculated using a community detection algorithm (Leicht and Newman 2008). We ranked networks according to each macro-scale index (connectance, nestedness, and modularity) in turn and divided networks into subsets of 50 according to this ranking. For example, when ranking networks by connectance there would be 400 subsets, each containing 50 networks with similar values of connectance.

For each subset, we calculated the multivariate distance between each network and the subset centroid representing the ‘typical’ structure for each subset. Specifically, we used a distance measure based on the correlation between vectors describing network structure. Distances were calculated using the ‘betadisper’ function from the ‘vegan’ R package (R Core Team 2015, Oksanen et al. 2016). We did this at both the macro- and meso-scale. Macro-scale vectors included the number of species in the first set, the number of species in the second set, and two of connectance, nestedness, and modularity (having excluded the metric used for ranking networks). For example, if connectance was used as the ranking property, the macro-scale structure vector would include the number of species in the first set, the number of species in the second set, nestedness and modularity. The vectors describing meso-scale structure were the structural signatures of motif frequencies, 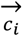, as described above. Correlations between vectors ranged between 1 and −1; to ensure non-negative values of distance, we subtracted each correlation value from 1. To control for the possibility that some subsets might have more variable structure than others, we then normalised distances by dividing by the maximum distance in each subset, giving values between 0 (for identical networks) and 1 (for completely different networks). We repeated this procedure using each of connectance, nestedness, and modularity as the ranking variable, to give three views of the variability of network structure at macro- and meso-scales. We then fitted generalised additive models (GAMs) relating variability in network structure to the ranking variable and to the scale at which variation was measured, using macro-scale variation as the intercept level. We fitted separate smoothed slopes for each scale. All GAMs were fitted using the ‘gam’ function from the ‘mgcv’ R package (Wood 2011, R Core Team 2015).

At the macro-scale, the mean variations when ranked by connectance, nestedness and modularity were 0.0399 (se ± 0.0010), 0.1261 (se ± 0.0015) and 0.0632 (se ± 0.0019), respectively. The corresponding mean variations at the meso-scale were 0.1347 (se ± 0.0029), 0.1232 (se ± 0.0027) and 0.1248 (se ± 0.0028). These results show that, for a given level of connectance or modularity, networks that appear similar at the macro-scale can be composed of very different indirect interaction structures: meso-scale structural signatures based on motifs generally showed much more dissimilarity than macro-scale measures of structure. Specifically, for connectance and modularity respectively, the motif framework captured 3.38 and 1.97 times more variation in indirect interactions than traditional whole-network indices. This interpretation is confirmed by the GAMs, where meso-scale variation was significantly greater than macro-scale variation (meso-scale intercepts were 0.095 and 0.062 for connectance and modularity, respectively, with p < 0.001 in both cases; Fig. 5a-b). When networks were ordered by nestedness, the picture was more complicated: when mean nestedness was low, meso-scale variation was higher than macro-scale variation, but when mean nestedness was high this trend was reversed (Fig. 5c). We note, however, that the standard error about the mean variation when networks were ranked by nestedness was twice as large for meso-scale structure than macro-scale. Overall, therefore, the increased variation (or increased variability in variation) in indirect interactions, highlights the problem of describing network structure by its macro-scale properties alone.

**Figure 5:**
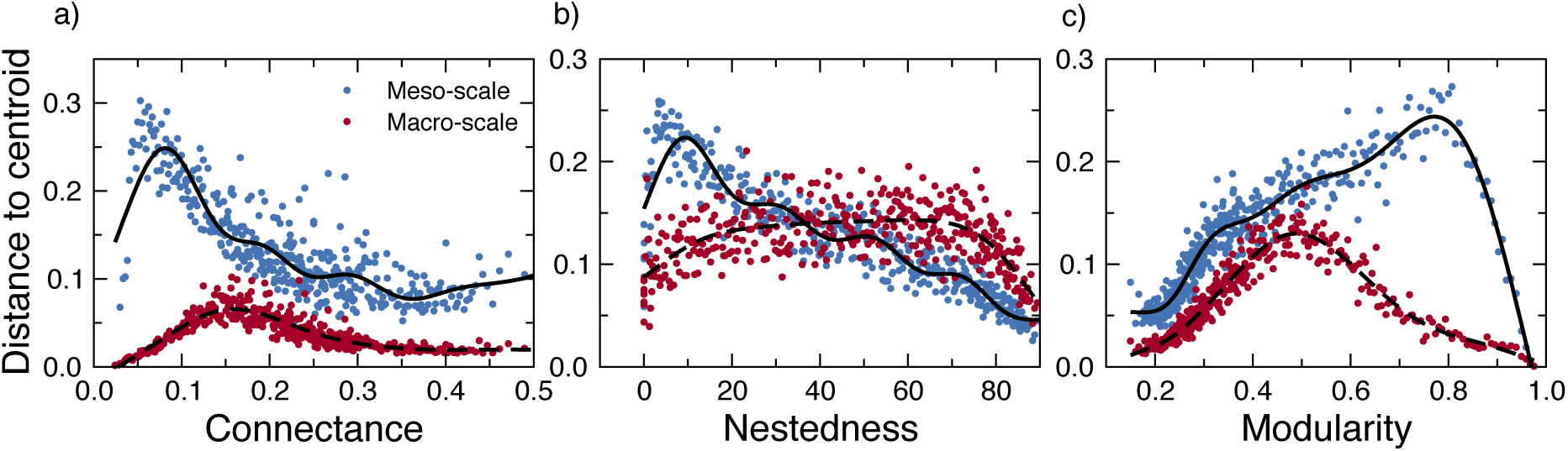
Network variation (normalised mean distance to group centroid) against mean connectance (left), nestedness (centre) and modularity (right) for all networks. Points represent subsets of networks.

## Robustness to sampling effort

A major challenge among studies of ecological networks is that completely sampling a web of interactions is difficult: both species and their interactions can be missed, especially if they are rare or hard to detect (Jordano 2016). Many network indices are sensitive to sampling effects (Dorado et al. 2011, Rivera-Hutinel et al. 2012, Fründ et al. 2016), though nestedness may be relatively robust (Nielsen and Bascompte 2007). To assess the sensitivity of the motif framework to sampling biases, we simulated different levels of sampling effort on 26 empirical, quantitative pollination and seed dispersal networks obtained from the Web of Life repository (www.web-of-life.es; Supplementary Table 1). In field studies, plant-animal interaction networks are usually sampled by observing plants and recording the animals that visit them (Jordano 2016). To replicate this process *in silico*, we sampled networks in two stages (de Aguiar et al. 2017). First, we sampled a proportion, *p*, of plant species to simulate the likely scenario that not all plant species are observed when surveying a site (Jordano 2016). Species with more partners had a higher chance of being sampled, as generalist species tend to be more abundant (Fort *et al.* 2016; though see Supplementary Fig. 1 for results where species had a random probability of being selected). Second, for each selected plant species, we sampled a proportion, *q*, of their interactions to simulate the fact that not all interactions are observed (Dormann et al. 2009, Poisot et al. 2012); stronger interactions, indicating more frequent visits between plants and animals, had a higher probability of being sampled (de Aguiar et al. 2017). We repeated this process for different values of *p* and *q* between 0.5 and 1, performing 1000 randomisations at each *p*-*q* combination. We decomposed each sampled network into its constituent motifs and recorded each network’s motif structural signature and the motif role signatures of each species. We then measured R^2^ between the network structural signature or species role signature of the sampled network and those of the corresponding ‘true’ network containing all species and interactions. Further details of the simulations are given in Supplementary Methods. We found that both network structural signatures and species role signatures were remarkably robust to sampling effects. Even when only 50% of plant species and 50% of their interactions were sampled, the mean R^2^ between the sampled and ‘true’ network signatures was 0.91 (Fig. 6a). At this same level of sampling, the mean R^2^ between sampled and ‘true’ species role signatures was 0.90 (Fig. 6b). That motifs appear robust to sampling effects is encouraging for future studies adopting this framework.

**Figure 6:**
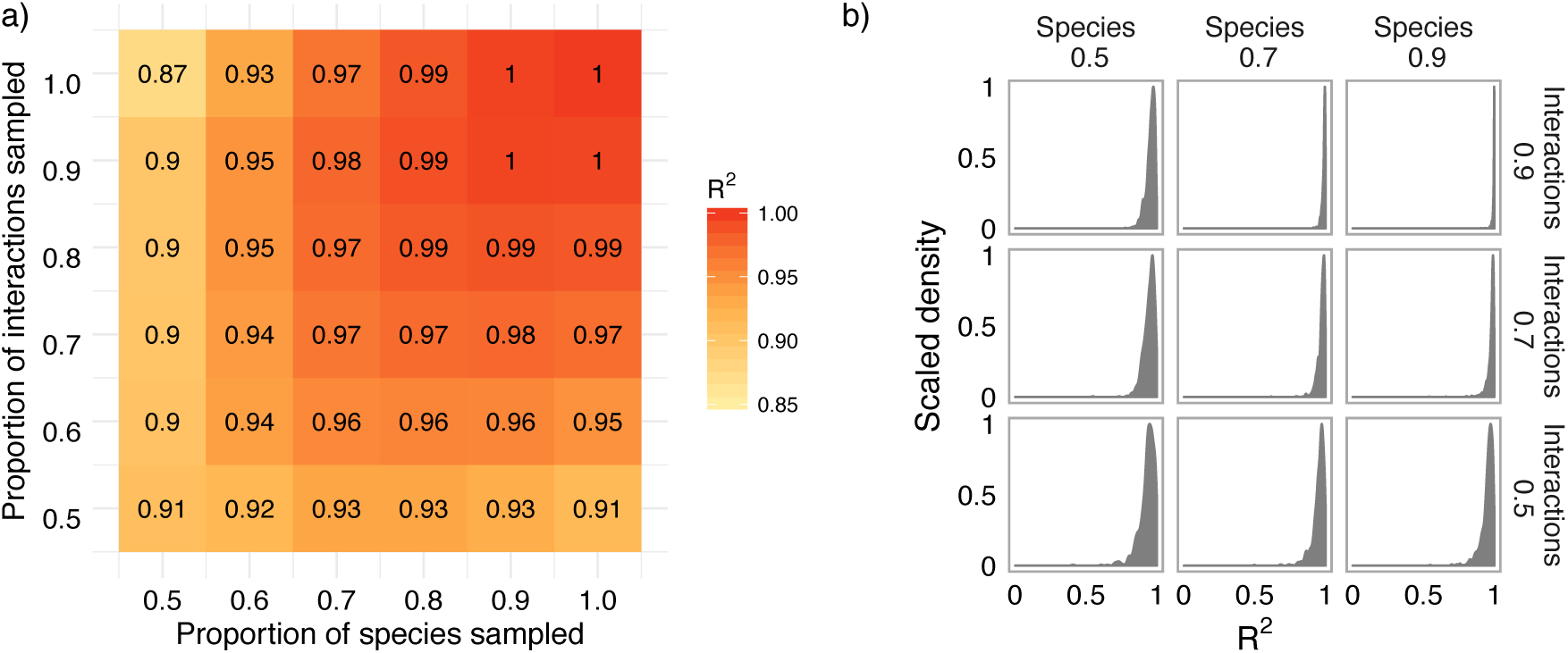
Results of simulations assessing the sensitivity of the motif framework to variation in sampling effort. (a) The mean R^2^ between the network structural signatures of the sampled networks and the structural signatures of their corresponding ‘true’ networks across all 26 networks for different levels of species and interaction removal. (b) Distribution of mean R^2^ between species role signatures in sampled networks and species role signatures in their corresponding ‘true’ networks for all species across all 26 networks for different levels of species and interaction removal. Note that the value in each cell in (a) is not necessarily the mean R^2^ across all 26 networks: sampled versions of some smaller networks were too small or disconnected to contain larger motifs, preventing the calculation of a structural signature which was comparable to that of the ‘true’ network. These networks were therefore excluded when calculating the mean R^2^. The number of webs in each cell is given in Supplementary Fig. 2. Similarly, in (b) each panel does not contain the R^2^ of all 1008 species as some species in smaller networks did not appear in larger motifs, particularly at high levels of species and interaction removal. The proportion of species in each panel is given in Supplementary Fig. 3.

## Indirect interactions in empirical plant-pollinator networks

Here we present a case study comparing the roles of two pollinator species over time. Data was from four mountaintop plant-pollinator communities in the Seychelles, sampled over the flowering season in eight consecutive months between September 2012 and April 2013 (Kaiser-Bunbury *et al.* 2017; Supplementary Table 2). Restoration by removal of exotic plants from these communities resulted in pollinator species becoming more generalised. This pattern was driven largely by two abundant, highly generalist pollinator species, one native (*Lasioglossum mahense*) and one non-native (*Apis mellifera*) (Kaiser-Bunbury et al. 2017). These two abundant, super-generalist species may have similar strategies for partner selection and therefore play similar roles in the community. This is the result found in the original study: both species had similar levels of specialisation (quantified using the specialisation index *d’*, which measures the extent to which species deviate from a random sampling of available partners (Blüthgen et al. 2006)): 0.17 ± 0.10 and 0.22 ± 0.18 for *Lasioglossum mahense* and *Apis mellifera* respectively. Alternatively, two abundant, super-generalists could minimise competition by exploiting different areas of ‘interaction niche space’ and therefore have different roles. To test these alternatives, we calculated the motif role signatures of both species at each site in each monthly network, giving a detailed view of how each species is embedded in the community over time. We used permutational multivariate analysis of variance (PERMANOVA), stratified by site, to assess if there are significant differences between the roles the two species play in the four communities. PERMANOVA is similar to ANOVA but compares multivariate differences within and between groups without assuming normality or Euclidean distances (Anderson 2001). We used Bray-Curtis distance as the dissimilarity measure, as it is suitable for a variety of ecological data, including motifs (Faith et al. 1987, Anderson and Robinson 2003, Baker et al. 2015). PERMANOVAs were run with 10000 permutations.

The PERMANOVA analysis showed that *Lasioglossum mahense* and *Apis mellifera* had significantly different roles over time (*F*_1,62_, p = 0.0496), exploiting different areas of interaction niche space. This means that, while Kaiser-Bunbury *et al*. (2017)used the species-level metric *d’* to show that both species were super-generalists, considering indirect interactions reveals that they are generalist in different ways. This result is visualised in Fig. 7. More positive values of the first NMDS axis are associated with motif positions where more specialist pollinators compete with generalist pollinators for a shared plant resource, while negative values are associated with positions where generalist pollinators visit specialist plants with little competition. More positive values of the second NMDS axis are associated with positions where pollinators visit plants which are also visited by generalist species; negative values are associated with positions where pollinators visit plants which are also visited by specialist species. *Lassioglossum mahense* generally occupies higher values of both NMDS axes than *Apis mellifera*. Therefore, while both species are generalists, *Lassioglossum mahense* is in greater competition with generalist pollinators than *Apis mellifera* which visits more specialist plants and competes with more specialist pollinators. These differences in indirect interactions are essential for understanding the ecology of these two species and are missed using the *d’* index alone. All PERMANOVA tests and NMDS analyses were conducted in the R package vegan (Oksanen et al. 2016).

**Figure 7:**
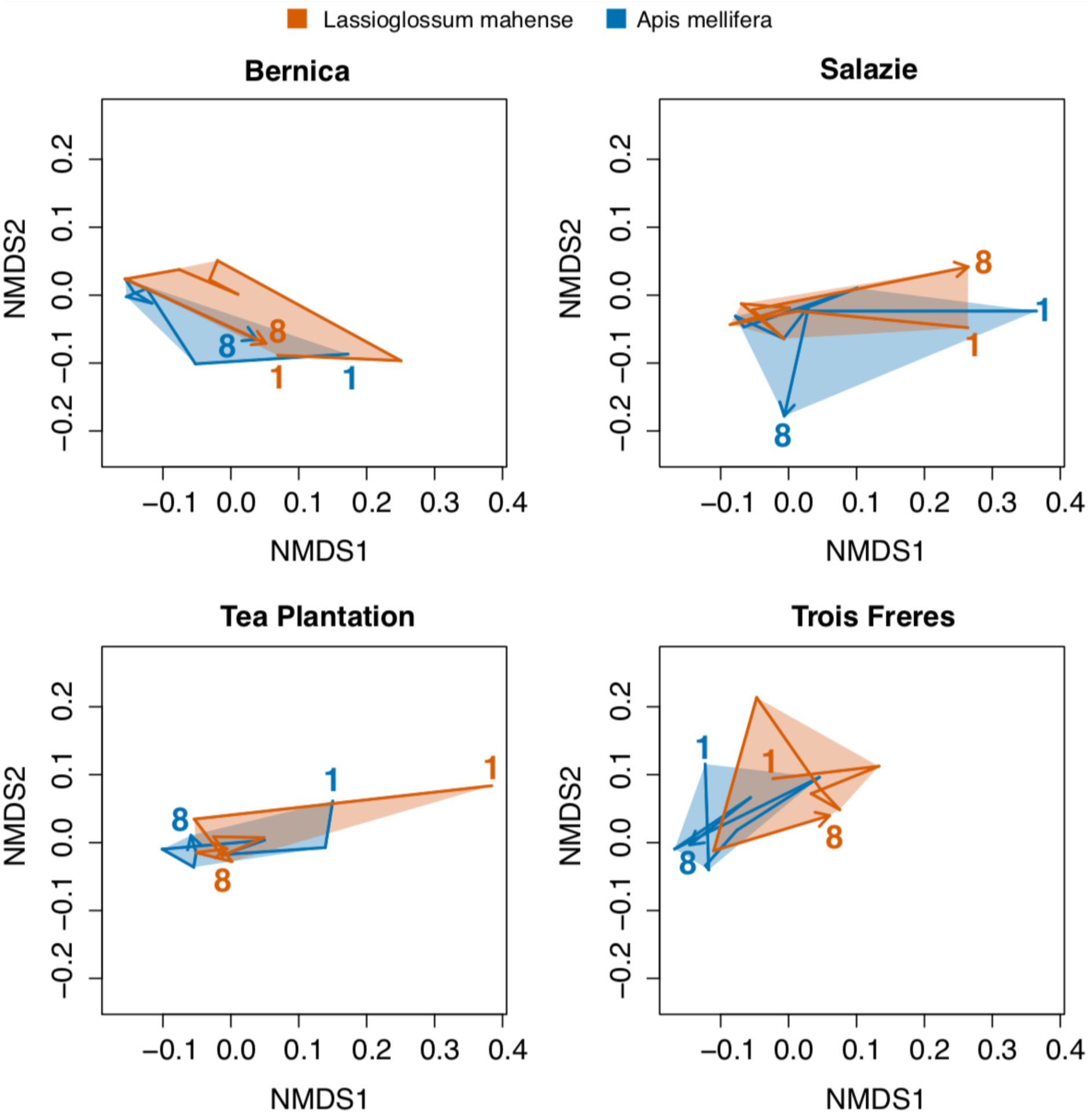
The movement of Lasioglossum mahense and Apis mellifera through interaction niche space over eight months in four sites (Bernica, Salazie, Tea Plantation and Trois Freres). Each vertex represents the role of a species in a monthly network. Numbers ‘1’ and ‘8’ indicate the first and last sampling month, respectively. Shaded polygons are convex hulls containing the vertices of each species.

## Potential applications

Characterising the structure of species interaction networks is a key component of many areas of ecological research, such as robustness to extinctions (Kaiser-Bunbury et al. 2010), ecosystem functioning (Coux et al. 2016) and macroecology (Araújo and Luoto 2007, Staniczenko et al. 2017). It is essential to incorporate indirect interactions into all these analyses, suggesting that the framework presented here has wide applicability to a diverse range of topics, systems and interaction types. In particular, we suggest the motif framework may be beneficial for studies where the scale of interest is at the species level, such as examining how invasive species integrate into communities (Vilà et al. 2009, Stouffer et al. 2014); or when within-network phenomena are the focus, such as studies of rewiring and network variability over time (Olesen et al. 2008, Kaiser-Bunbury et al. 2010). As indirect interactions are likely to be of increased importance when investigating these types of questions, we strongly caution against using only conventional indices which mask these interactions. Adopting multidimensional descriptions of network structure as collections of their component parts also opens up new ways to answer a diverse range of questions such as those concerning competitive exclusion, species packing and functional redundancy (Blonder et al. 2014). For example, does interaction distinctiveness correlate with functional distinctiveness? Do species have overlapping or disjoint roles? Do indirect interaction structures vary over space and time?

We have shown how easy it is for similar-looking networks to be composed of very dissimilar parts. We therefore expect that much valuable information on indirect interactions has been ignored, intentionally or unintentionally. This realisation yields a series of hypotheses about how our understanding of bipartite ecological networks may change if indirect interactions were incorporated. For example, uncovering indirect interactions could revise our understanding of how invariant network structure is across space and time. Several studies have shown that network structure is relatively stable in the presence of temporal and spatial turnover in species and interaction identity (Petanidou et al. 2008, Dáttilo et al. 2013). However, these studies have considered only global descriptors of structure that likely mask meso-scale structural variation in indirect interactions. We anticipate that, if indirect interactions were considered, network structure may not be as invariant to compositional turnover as previously identified.

We also hypothesise that incorporating indirect interactions may improve predictions of network structure and our understanding of the mechanisms underpinning network assembly. Understanding the processes that govern the formation of species interactions is essential for predicting the structure of novel communities under global changes (Eklöf et al. 2013). However, current attempts often involve assessing how well different mechanisms, such as neutral effects, morphological matching and phenological overlap, predict network- or species-level indices (Vázquez et al. 2009, Verdú and Valiente-Banuet 2011, Sayago et al. 2013, Vizentin-Bugoni et al. 2014). For example, Vázquez et al. (2009)show that data on abundance and phenology can accurately predict macro-scale indices such as connectance and nestedness. This approach is problematic because many indices are insensitive to changes in network topology. Models can therefore accurately predict index values while incorrectly predicting pairwise interactions (Fox 2006, Olito and Fox 2015). Such models may be of limited utility in helping to understand the processes underlying network structure. To improve models, structural signatures based on motifs could be used instead as a benchmark of predictive performance. As the motif framework is much more sensitive to changes in network topology than indices are, it would be harder for models to accurately predict a structural signature while incorrectly predicting pairwise interactions. We therefore expect that adopting the motif framework could change both our understanding of the processes governing interactions and improve our ability to predict novel communities.

Finally, incorporating indirect interactions could improve understanding of the functional consequences of community structure (Thompson et al. 2012, Poisot et al. 2013). For example, pollinators with more distinct traits (traits furthest from the community average) tend to have fewer interaction partners (Coux et al. 2016). One hypothesis for this pattern is a trade-off between reducing competition with other pollinators by having original traits and needing to retain interaction partners (Vamosi et al. 2014, Coux et al. 2016). The motif framework could explicitly test this hypothesis by assessing whether functionally original species appear primarily in motif positions where there is low competition between pollinators.

## Limitations and challenges

Currently, bipartite motifs are only defined for qualitative networks, where interactions are present or absent. This contrasts with quantitative networks where interactions are weighted in proportion to their relative strength. Using only qualitative information, rare species or interactions can exert a disproportionate influence on network metrics (Banašek-Richter et al. 2004). The loss of detail on indirect interactions resulting from the use of conventional indices is, however, likely to be equal to or greater than the loss of information resulting from using qualitative instead of quantitative networks. As shown in the example above, *d’*, an index which uses quantitative information on interaction weights, could not distinguish the roles of *Lasioglossum mahense* and *Apis mellifera*, while qualitative motifs could. We also note that qualitative versions of many conventional metrics (such as connectance and degree) are frequently used to characterise quantitative networks instead of their weighted counterparts. While methods to enumerate weighted motifs will likely be developed in the future, there are already a number of tractable methods to incorporate quantitative information in motif analyses. For example, interactions within motifs can be classified as ‘strong’ or ‘weak’ depending on whether a given interaction’s strength is greater or lesser, respectively, than the median strength (Rodríguez-Rodríguez et al. 2017). Alternatively, a suite of qualitative networks can be assembled by sampling a quantitative network in proportion to the strength of each interaction (Baker et al. 2015). This creates an ensemble of qualitative resampled networks where stronger interactions appear more frequently than weaker ones. Analyses can then be repeated using each of the resampled networks as input. This creates a distribution of p-values or effect sizes associated with a particular analysis, which can then be compared to the results obtained using a binary version of the original quantitative network.

Finally, it is important to note that, like indices, the motif framework also results in a loss of information when characterising network structure: transforming a network into a structural signature or ensemble of species’ role signatures is unique, while the reverse is not. Some loss of information is inevitable so long as we must summarize networks in order to analyse them. However, motifs are substantially less interaction inelastic than indices, and therein lies their advantage.

## Concluding remarks

Indirect interactions are a widespread and important component of ecological communities, essential for understanding species roles and the structure of biotic interactions. However, to date the dominant paradigm has been to describe community structure using a wide variety of indices that often ignore indirect interactions. Here we have presented a framework that moves beyond such metrics to conceptualise networks as a series of component building blocks or ‘motifs’. By thinking of networks in this way, we have shown that indirect interactions can be identified and quantified. We do not advocate widespread abandonment of indices, but instead aim to raise awareness of their significant limitations. We hope that motifs will exist alongside indices to form the basis of a new paradigm among studies of bipartite ecological networks. Given the increasingly large amount of ecological network data available, and the rapid growth in computational capacity to analyse these data, there is now a timely opportunity to make motifs a standard part of the analytical toolkit for studying bipartite systems. Such an approach could enable novel perspectives and insights into the ecology and evolution of many important communities.

## Data accessibility

All analyses in this manuscript use open access data which are already archived in public repositories.

## Acknowledgements

Funding – BIS is supported by the Natural Environment Research Council as part of the Cambridge Earth System Science NERC DTP [NE/L002507/1]. ARC, NJB, and DBS were all supported by Marsden Fund Fast-Start grant UOC-1101, administered by the Royal Society of New Zealand. NJB was also supported by a BlueFern HPC PhD scholarship and DBS by a Rutherford Discovery Fellowship, also administered by the RSNZ. ARC was also supported by a Formas grant (#942-2015-1262, awarded to Anna Eklöf). LVD was supported by the Natural Environment Research Council (grants NE/K015419/1 and NE/N014472/1). WJS is funded by Arcadia.

## Author contributions

BIS conceived the study, performed analyses and wrote the first draft of the manuscript. BIS, ARC and DBS designed analyses and interpreted results. ARC and NJB performed the analysis demonstrating meso-scale dissimilarity in networks with similar macro-scale properties. All authors contributed to writing the manuscript.

## References

Almeida-Neto, M. et al. 2008. A consistent metric for nestedness analysis in ecological systems: Reconciling concept and measurement. - Oikos 117: 1227–1239.

Anderson, M. J. 2001. A new method for non parametric multivariate analysis of variance. - Austral Ecol. 26: 32–46.

Anderson, M. J. and Robinson, J. 2003. Generalized discriminant analysis based on distances. - Aust. New Zeal. J. Stat. 45: 301–318.

Araújo, M. B. and Luoto, M. 2007. The importance of biotic interactions for modelling species distributions under climate change. - Glob. Ecol. Biogeogr. 16: 743–753.

Bailey, J. K. and Whitham, T. G. 2007. Biodiversity is related to indirect interactions among species of large effect. - In: Ohgushi, T. et al. (eds), Ecological communities: plant mediation in indirect interaction webs. Cambridge University Press, pp. 306–328.

Baker, N. J. et al. 2015. Species’ roles in food webs show fidelity across a highly variable oak forest. - Ecography (Cop.). 38: 130–139.

Banašek-Richter, C. et al. 2004. Sampling effects and the robustness of quantitative and qualitative food-web descriptors. - J. Theor. Biol. 226: 23–32.

Blonder, B. et al. 2014. The n-dimensional hypervolume. - Glob. Ecol. Biogeogr. 23: 595–609.

Blüthgen, N. et al. 2006. Measuring specialization in species interaction networks. - BMC Ecol. 6: 9.

Carvalheiro, L. G. et al. 2014. The potential for indirect effects between co-flowering plants via shared pollinators depends on resource abundance, accessibility and relatedness. - Ecol. Lett. 17: 1389–1399.

Cirtwill, A. R. and Stouffer, D. B. 2015. Concomitant predation on parasites is highly variable but constrains the ways in which parasites contribute to food web structure. - J. Anim. Ecol. 84: 734–744.

Cirtwill, A. R. et al. 2018. Between-year changes in community composition shape species’ roles in an Arctic plant--pollinator network. - Oikos in press.

Coux, C. et al. 2016. Linking species functional roles to their network roles. - Ecol. Lett. 19: 762–770.

Dáttilo, W. et al. 2013. Spatial structure of ant-plant mutualistic networks. - Oikos 122: 1643–1648.

de Aguiar, M. A. M. et al. 2017. Revealing biases in the sampling of ecological interaction networks. - arXiv: 1708.01242.

Dorado, J. et al. 2011. Rareness and specialization in plant-pollinator networks. - Ecology 92: 19–25.

Dormann, C. F. et al. 2009. Indices, Graphs and Null Models: Analyzing Bipartite Ecological Networks. - Open Ecol. J. 2: 7–24.

Dupont, Y. L. et al. 2014. Spatial structure of an individual-based plant-pollinator network. - Oikos 123: 1301–1310.

Eklöf, A. et al. 2013. The dimensionality of ecological networks. - Ecol. Lett. 16: 577–583.

Faith, D. P. et al. 1987. Compositional dissimilarity as a robust measure of ecological distance. - Vegetatio 69: 57–68.

Fontaine, C. et al. 2011. The ecological and evolutionary implications of merging different types of networks. - Ecol. Lett. 14: 1170–1181.

Fort, H. et al. 2016. Abundance and generalisation in mutualistic networks: Solving the chicken- and-egg dilemma. - Ecol. Lett. 19: 4–11.

Fox, J. W. 2006. Current food web models cannot explain the overall topological structure of observed food webs. - Oikos 115: 97–109.

Frank van Veen, F. J. et al. 2006. Apparent competition, quantitative food webs, and the structure of phytophagous insect communities. - Annu. Rev. Entomol. 51: 187–208.

Fründ, J. et al. 2016. Sampling bias is a challenge for quantifying specialization and network structure: Lessons from a quantitative niche model. - Oikos 125: 502–513.

Guimarães, P. R. et al. 2017. Indirect effects drive coevolution in mutualistic networks. - Nature 550: 511–514.

Jordán, F. et al. 2007. Quantifying positional importance in food webs: A comparison of centrality indices. - Ecol. Modell. 205: 270–275.

Jordano, P. 2016. Sampling networks of ecological interactions. - Funct. Ecol. 30: 1883–1893.

Kaiser-Bunbury, C. N. et al. 2010. The robustness of pollination networks to the loss of species and interactions: A quantitative approach incorporating pollinator behaviour. - Ecol. Lett. 13: 442–452.

Kaiser-Bunbury, C. N. et al. 2017. Ecosystem restoration strengthens pollination network resilience and function. - Nature 542: 223–227.

Kashtan, N. et al. 2004. Topological generalizations of network motifs. - Phys. Rev. E - Stat. Physics, Plasmas, Fluids, Relat. Interdiscip. Top. 70: 12.

Leicht, E. A. and Newman, M. E. J. 2008. Community structure in directed networks. - Phys. Rev. Lett. 100: 118703.

Martínez, D. et al. 2014. Consistency and reciprocity of indirect interactions between tree species mediated by frugivorous birds. - Oikos 123: 414–422.

Menke, S. et al. 2012. Plant-frugivore networks are less specialized and more robust at forest-farmland edges than in the interior of a tropical forest. - Oikos 121: 1553–1566.

Milo, R. et al. 2002. Network motifs: simple building blocks of complex networks. - Science (80-.). 298: 824–827.

Mitchell, R. J. et al. 2009. New frontiers in competition for pollination. - Ann. Bot. 103: 1403–1413.

Morales, C. L. and Traveset, A. 2009. A meta-analysis of impacts of alien vs. native plants on pollinator visitation and reproductive success of co-flowering native plants. - Ecol. Lett. 12: 716–728.

Morris, R. J. et al. 2004. Experimental evidence for apparent competition in a tropical forest food web. - Nature 428: 310–313.

Morris, R. J. et al. 2014. Antagonistic interaction networks are structured independently of latitude and host guild. - Ecol. Lett. 17: 340–349.

Nielsen, A. and Bascompte, J. 2007. Ecological networks, nestedness and sampling effort. - J. Ecol. 95: 1134–1141.

Oksanen, J. et al. 2016. vegan: Community Ecology Package. R package version 2.4-0. in press.

Olesen, J. M. et al. 2008. Temporal dynamics in a pollination network. - Ecology 89: 1573–1582.

Olito, C. and Fox, J. W. 2015. Species traits and abundances predict metrics of plant-pollinator network structure, but not pairwise interactions. - Oikos 124: 428–436.

Petanidou, T. et al. 2008. Long-term observation of a pollination network: Fluctuation in species and interactions, relative invariance of network structure and implications for estimates of specialization. - Ecol. Lett. 11: 564–575.

Poisot, T. et al. 2012. A comparative study of ecological specialization estimators. - Methods Ecol. Evol. 3: 537–544.

Poisot, T. et al. 2013. Trophic complementarity drives the biodiversity-ecosystem functioning relationship in food webs. - Ecol. Lett. 16: 853–861.

R Core Team 2015. R: A language and environment for statistical computing. in press.

Rivera-Hutinel, A. et al. 2012. Effects of sampling completeness on the structure of plant-pollinator networks. - Ecology 93: 1593–1603.

Rodríguez-Rodríguez, M. C. et al. 2017. Functional consequences of plant-animal interactions along the mutualism-antagonism gradient. - Ecology 98: 1266–1276.

Saavedra, S. et al. 2009. A simple model of bipartite cooperation for ecological and organizational networks. - Nature 457: 463–466.

Saracco, F. et al. 2017. Inferring monopartite projections of bipartite networks: An entropy-based approach. - New J. Phys. 19: 53022.

Sayago, R. et al. 2013. Evaluating factors that predict the structure of a commensalistic epiphyte-phorophyte network. - Proc. R. Soc. B Biol. Sci. 280: 20122821–20122821.

Staniczenko, P. P. A. et al. 2017. Linking macroecology and community ecology: refining predictions of species distributions using biotic interaction networks. - Ecol. Lett. 20: 693–707.

Stouffer, D. B. et al. 2012. Evolutionary Conservation of Species’ Roles in Food Webs. - Science (80-.). 335: 1489–1492.

Stouffer, D. B. et al. 2014. How exotic plants integrate into pollination networks. - J. Ecol. 102: 1442–1450.

Strauss, S. Y. 1991. Indirect Effects in Community Ecology - Their Definition, Study and Importance. - Trends Ecol. Evol. 6: 206–210.

Tack, A. J. M. et al. 2011. Can we predict indirect interactions from quantitative food webs? - An experimental approach. - J. Anim. Ecol. 80: 108–118.

Thompson, R. M. et al. 2012. Food webs: Reconciling the structure and function of biodiversity. - Trends Ecol. Evol. 27: 689–697.

Vamosi, J. C. et al. 2014. Pollinators visit related plant species across 29 plant-pollinator networks. - Ecol. Evol. 4: 2303–2315.

Vandermeer, J. et al. 1985. Indirect facilitation and mutualism. - In: Boucher, D. H. (ed), The Biology of Mutualism: Ecology and Evolution. Oxford University Press, pp. 326–343.

Vázquez, D. P. et al. 2009. Evaluating multiple determinants of the structure of plant-animal mutualistic networks. - Ecology 90: 2039–2046.

Verdú, M. and Valiente-Banuet, A. 2011. The relative contribution of abundance and phylogeny to the structure of plant facilitation networks. - Oikos 120: 1351–1356.

Vilà, M. et al. 2009. Invasive plant integration into native plant-pollinator networks across Europe. - Proc. R. Soc. B Biol. Sci. 276: 3887–3893.

Vizentin-Bugoni, J. et al. 2014. Processes entangling interactions in communities: forbidden links are more important than abundance in a hummingbird-plant network. - Proc. R. Soc. B Biol. Sci. 281: 20132397.

Wood, S. N. 2011. Fast stable restricted maximum likelihood and marginal likelihood estimation of semiparametric generalized linear models. - J. R. Stat. Soc. Ser. B Stat. Methodol. 73: 3–36.

Wootton, J. T. 2002. Indirect effects in complex ecosystems: recent progress and future challenges. - J. Sea Res. 48: 157–172.

Zhou, T. et al. 2007. Bipartite network projection and personal recommendation. - Phys. Rev. E - Stat. Nonlinear, Soft Matter Phys. 76: 46115.

